# Modulating long-range energetics via helix stabilization: a case study using T4 lysozyme

**DOI:** 10.1101/353649

**Authors:** Sabriya N. Rosemond, Kambiz M. Hamadani, Jamie H.D. Cate, Susan Marqusee

## Abstract

Cooperative protein folding requires distant regions of a protein to interact and provide mutual stabilization. The mechanism of this long-distance coupling remains poorly understood. Here, we use T4 lysozyme (T4L*) as a model to investigate long-range communications across a globular protein. T4L* is composed of two structurally distinct subdomains, although it behaves in a two-state manner at equilibrium. The subdomains of T4L* are connected via two topological connections: the N-terminal helix that is structurally part of the C-terminal subdomain (the A-helix) and a long helix that spans both subdomains (the C-helix). To understand the role that the C-helix plays in cooperative folding, we analyzed a circularly permuted version of T4L* (CP13*), whose subdomains are connected only by the C-helix. We demonstrate that when isolated as individual fragments, both subdomains of CP13* can fold autonomously into marginally stable conformations. The energetics of the N-terminal subdomain depend on the formation of a salt bridge known to be important for stability in the full-length protein. We show that the energetic contribution of the salt bridge to the stability of the N-terminal fragment increases when the C-helix is stabilized, such as occurs upon folding of the C-terminal subdomain. These results suggest a model where long-range energetic coupling is mediated by helix stabilization.

---

In this work, we investigate how a helix spanning the two subdomains of T4 lysozyme* couples distant regions of the protein. We find evidence for a model of long-distance coupling that relies on the cooperative nature of helix formation to stabilize a long-range tertiary salt bridge interaction at one end of the helix and thereby couple the folding of T4 lysozyme’s subdomains. This mechanistic model may have implications for co-translational folding.

## Introduction

Cooperativity is a hallmark of globular proteins. At equilibrium, many small (<200 residues) globular proteins fold in an apparent two-state manner (U⇌N), populating either a completely folded or unfolded conformation^1,2^. The stability of the folded protein (ΔG_UN_ = G_N_−G_U_) is the sum of many interactions, both local and long-range. While the chemical nature and mechanism of the short-range, local interactions can be easy to identify and investigate, it is difficult to probe the long-range interactions that cause the structure and energetics in one region to be coupled to those in another region and thus lead to two-state behavior. For repeat proteins, such as ankrin domains, this coupling has been attributed to a large interfacial energy that, in some cases, overshadows the lack of intrinsic stability of the individual repeats^3,4^.

Here, we explore coupling between distant regions of a protein using the protein T4 lysozyme*, T4L* (* denotes the cysteine-free pseudo-wild type variant^5^). T4L* is a well-studied globular protein; the stabilities and structures of hundreds of single-site mutants have provided a wealth of information about the role of side-chain interactions and their effect on protein stability^6^. The protein is composed of two distinct subdomains (Fig. 1A): the N-terminal subdomain (residues 13-74) and the C-terminal subdomain (residues 75-164 and 1-12) ^7,8^. The full-length protein unfolds in an apparent two-state manner with no notable populated intermediates (ΔG_unf_ ~14 kcal/mol)^8,9^. In isolation, however, the C-terminal subdomain folds autonomously into a marginally stable structure (2.1 kcal/mol), while the isolated N-terminal subdomain has been reported to be predominately unfolded^8^. Together, these results indicate that in the context of the full-length protein, folding of the domains is strongly coupled.

**Figure 1.**
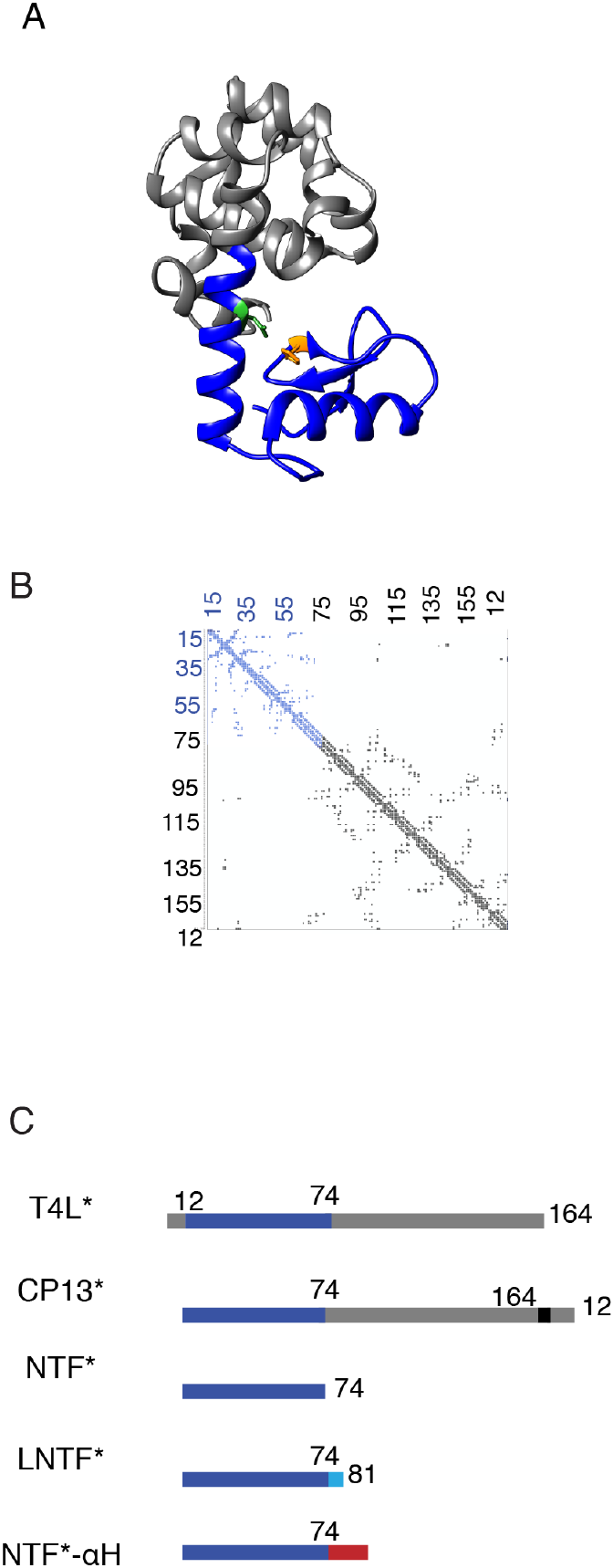
CP13* structure, subdomain architecture and constructs. A. Ribbon diagram of CP13*. N-terminal subdomain (dark blue) (PDB ID 2O4W) and C-terminal subdomain (gray) are connected by the long central C-helix. Residues involved in salt bridge are colored His31(orange) Asp70 (green). B. Contact map of CPxs13*. Interresidue contacts in the N-terminal subdomain in blue. Interresidue contacts in C-terminal subdomain in grey. C. Primary sequence representation of T4L*, CP13*, NTF*, LNTF*, and NTF*-αH. N-terminal subdomain in dark blue, C-terminal subdomain in grey. Extended N-terminal subdomain fragments LNTF (with residues 74-81 in light blue) and NTF-αH with alanine helix residues in red.

The nature of the strong coupling between the two subdomains is difficult to discern as almost all of the side-chain contacts in T4L* are within the individual subdomains^7,8^ (Fig. 1B). Thus, communication between the subdomains must derive from the topology or backbone connections between the two. There are two such connections: one at the end of the A-helix and one within the C-helix (Fig. 1). The A-helix (residues 1-12) is structurally part of the C-terminal subdomain, making this subdomain discontinuous in sequence^8^ and connects the two subdomains between residues 12 and 13. The C-helix is the central helix that spans both subdomains (residues 59-81, Fig. 1 A).

The role of this discontinuous subdomain architecture (the placement of helix A) was explored using a circular permutant, CP13*^8–11^ (Fig. 1A). CP13* begins at residue 13 with the A-helix (residues 1-12) appended to the C-terminus via a short flexible linker^8^. Thus, in CP13*, the C-terminal subdomain is contiguous in sequence. In spite of this rearrangement, CP13* and T4L* have similar structures (RMSD<1.0Å)^10^ and CP13* retains two-state behavior at equilibrium^8–10^. Thus, the discontinuous subdomain architecture of T4L* is not fully responsible for the coupling between the subdomains. It should be noted, however, that this permutation does have a small effect on the coupling between the two subdomains as determined by native-state hydrogen exchange (NSHX)^10^ and single molecule force spectroscopy^11^. NSHX studies on the full-length protein identified a rarely-populated, high-energy, partially-unfolded conformation under native conditions that consists of an extended folded C-terminal subdomain and an unfolded N-terminal subdomain^12^, and experiments with CP13* revealed an increase in the population of this intermediate^10^. Thus, the discontinuous subdomain architecture of T4L* aids in coupling the two regions, but it is not the whole story.

One of the most important side-chain interactions within T4L* is a buried salt bridge between His31 and Asp70 - a partially buried interaction within the N-terminal subdomain of the protein. This interaction stabilizes the native state of T4L* and CP13* by 3-5 kcal/mol^8,13^. In the present work, we identify an additional role for this salt bridge: coupling the two subdomains of T4L*. We revisit the isolated N-terminal subdomain and find that it is folding-competent in conditions that promote the formation of the H31-D70 salt bridge and its stability is dependent on this interaction. We also find that extending the N-terminal subdomain fragment to include the last seven residues of the C-helix, which are part of the C-terminal subdomain, stabilizes the interaction between these two residues and increases the stability of the N-terminal subdomain. This stabilizing effect can be recapitulated by adding an autonomously folding alanine-based helical sequence.

One of the partners in this salt bridge, residue 70, resides in the C-helix and is part of the N-terminal subdomain. We propose that helix stabilization provided by interactions at the other end of the C-helix (in the C-terminal subdomain) correctly orients D70 with respect to H31, thereby increasing its effective concentration and the salt-bridge stability. Thus, the C-helix-mediated coupling between the two subdomains appears to be transmitted via the helical backbone. This mechanism, coupling via stable helix formation, may be a general feature of long-range communication within proteins.

## Results

### The isolated N-terminal subdomain folds in a pH-dependent manner

A fragment encoding the N-terminal subdomain (residues 13-74 of T4L*, NTF*) was cloned, expressed and purified for biophysical studies (See Materials and Methods). Previous studies on a similar construct characterized the fragment as “predominantly unfolded”^8^, but we find that under slightly different experimental conditions (pH 5 versus 4.4), the fragment is folded as monitored by far-UV circular dichroism (CD) (Fig. 2A). The urea-induced equilibrium denaturation (monitored by the change in CD signal at 222nm, pH 5, 4°C) shows a sigmoidal transition, indicating cooperative unfolding (Fig. 2B). Although the transition is very broad, this denaturation curve can be fit to a two-state model^14^, yielding an extrapolated free energy (ΔG_unf_(H2O) of 1.60 ±0.13 kcal/mol and an m-value =1.13 ±0.01 kcal/mol*M (Fig. 2B and Table I). The m-value agrees with expectations based on the number of residues in the fragment (m-values have been found to correlate with buried surface area and size of a given polypeptide^15^), consistent with folding of the entire fragment.

**Figure 2.**
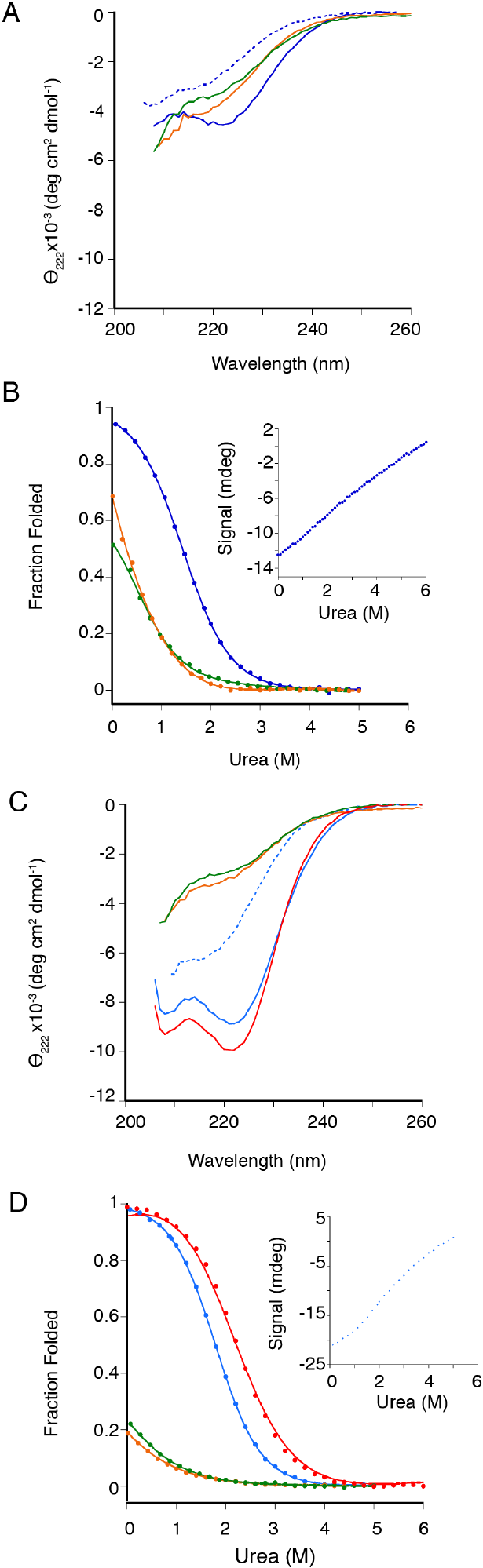
NTF* folds into a marginally stable structure and extending its sequence alters its structure and energetics. A. CD spectrum of NTF* at pH 5 (blue solid line) and pH 7 (blue dashed line) NTF* H31N (orange line) NTF* D70N (green line). B. Representative equilibrium denaturation curves of NTF* and variants monitored by CD at 222nm at pH 5, of NTF* (blue circle), NTF* H31N (orange circles) and NTF* D70N (green circles). Inset of NTF* urea denaturation melt at pH 7 (blue circles). C. CD spectra of LNTF* at pH 5 (light blue solid line) and pH 7 (light blue dashed line), LNTF* H31N at pH5 (orange line), LNTF* D70N at pH 5 (green line) and NTF*-αH at pH 5 (red). D. Representative equilibrium denaturation curves normalized to fraction folded. LNTF* (light blue circles) and NTF*-αH (red circles), LNTF* H31N (orange circles) and LNTF* D70N (green circles) at pH5. Inset of LNTF* urea-denaturation curve at pH 7 (light blue circles). All data taken at 4°C.

**Table I.**
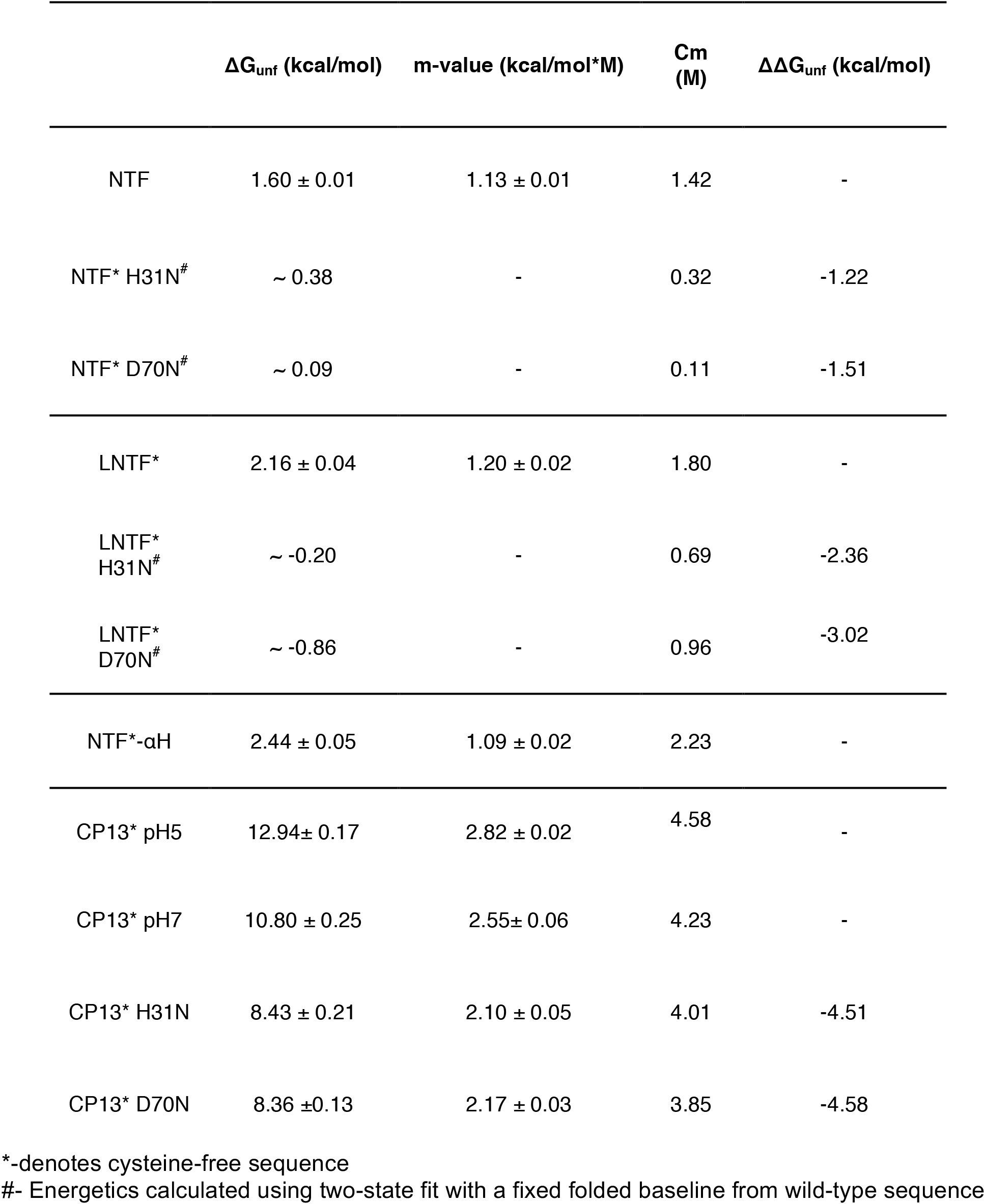
Energetics of fragment and full-length sequences derived from urea-denaturant melts

The structure and energetics of NTF* are pH-dependent. CD spectra of NTF* at pH 5 and pH 7 show a difference in magnitude and shape (Fig. 2A). At pH 7, the fragment is no longer folded, as judged by the loss of a cooperative unfolding transition with urea (Fig. 2B, inset). This pH dependence may be explained by the partially buried salt bridge (H31-D70) within the N-terminal subdomain^7,8,13^.

### H31 and D70 are important for the folding and stability of NTF*

To probe the role of these residues in NTF*, we generated two single-site variants, NTF* H31N and NTF* D70N. The CD spectra suggest that, unlike NTF*, both variants are predominately unfolded under native-like conditions (Fig. 2A). Urea-induced denaturation of these variants (pH 5, 4°C) monitored by CD at 222 nm shows no notable folded baseline, demonstrating that, even under conditions of no or low denaturant, the fragments are in the transition zone (Fig. 2B). To estimate how these mutations affect NTF*’s stability, we carried out a two-state fit on these data, fixing the folded baseline to the parameters from the wild-type NTF* fit. This analysis yields an estimate of ΔG_unf_ ~ 0.31 kcal/mol for NTF* H31N and ΔG_unf_ ~ 0.09 kcal/mol for NTF* D70N (Table I). The significant destabilization of NTF* that occurs upon mutating H31 and D70 suggests that these residues play a critical role in the stability of the isolated N-terminal subdomain.

### Extending the C-terminal helix enhances its stability

In full-length T4L*, the C-helix (residues 59-81) spans both subdomains: the majority of the helix is part of the N-terminal subdomain, but the final seven residues are in the C-terminal subdomain. To determine the effect of extending NTF* to include the complete C-helix, we generated a longer fragment encoding the N-terminal subdomain with the entire C-helix sequence, LNTF* (<u>L</u>ong-NTF*, residues 13-81). At pH 5, LNTF* is folded as monitored by far-UV CD and unfolds cooperatively as a function of urea (Fig. 2C and D). When fit with a two-state linear extrapolation model^14^, this unfolding curve results in a ΔG_unf_ = 2.16 ±0.04 kcal/mol with an m-value = 1.20 ±0.02 kcal/mol*M. Therefore, under these conditions, an extended N-terminal subdomain fragment is stabilized (0.56 ± .14 kcal/mol) compared to NTF*, which has the shorter C-helix (Table I). Similar to NTF*, the folding of LNTF* is pH-dependent, with less structure and a loss of cooperative unfolding at pH 7 compared to pH 5 (Fig. 2C and D, inset).

The same single-site variants investigated in the context of NTF* were also evaluated in the background of LNTF* (LNTF* H31N and LNTF* D70N). CD spectra and denaturation curves of these variants suggest that, like in the shorter fragment NTF*, they are predominately unfolded under native-like conditions. Using a similar approach as above to estimate the energetics of these variants, these curves were fit using a two-state model assuming the same baseline as LNTF*. Based on this analysis, the variants are significantly destabilized compared to LNTF* (LNTF* H31N: ΔG_unf_ ~ −0.20 kcal/mol; LNTF* D70N ΔGunf ~ −0.86 kcal/mol) (Table I, Fig. 2C and D).

To determine whether the increased stability of LNTF* as compared to NTF* is due to sequence-specific interactions or simply the addition of a helical segment, we appended a sequence known to form a stable helix^16^, (A(EAAAK)_3_A), to the C-terminus of NTF*, generating NTF*-aH. When monitored by CD, NTF*-αH is folded and has a cooperative urea-induced unfolding curve (Fig. 2D, Table I) (ΔG_unf_ = 2.44 ±0.05 kcal/mol, m-value = 1.09 ±0.02 kcal/mol*M (pH 5.0, 4°C)). The addition of this stable alpha helix increases NTF* stability by ~0.84 ±.14 kcal/mol at pH 5.

### Full-length CP13* populates an equilibrium intermediate in the absence of the H31-D70 salt bridge

The above fragment studies suggest that the H31-D70 salt bridge is critical to the stability of the isolated N-terminal subdomain (~1-2 kcal/mol). This same salt bridge is known to contribute 3-5 kcal/mol in full-length T4L*^13^. To probe the contribution of this interaction in full-length CP13*, we investigated CP13*’s stability as a function of pH and mutation. Equilibrium urea-denaturation studies analyzed with a two-state assumption showed a notable difference in the free energy and calculated m-value at pH 5 and 7 (pH 5: ΔG_unf_ = 12.94 ±0.17 kcal/mol, m-value= 2.82 ±0.02 kcal/mol*M; pH 7: ΔG_unf_ = 10.82 ±0.25 kcal/mol, m-value= 2.55± 0.06 kcal/mol*M, Fig. 3, Table I). The lower m-value at pH 7 suggests a breakdown in the two-state assumption due to a measureable population of an equilibrium intermediate^17–19^, which could indicate selective unfolding of the N-terminal subdomain at the higher pH.

**Figure 3.**
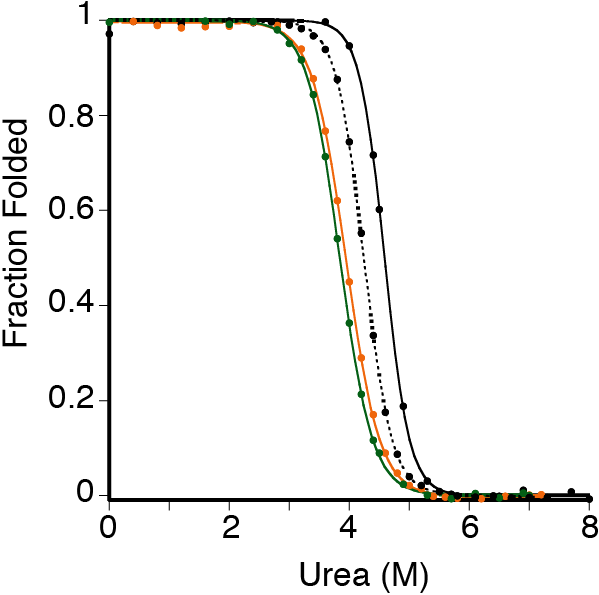
CP13* energetics are dependent on H31-D70 salt bridge. Representative denaturation melts of CP13* at pH5 (black solid line), pH7 (black dashed line), and CP13* H31N (orange line) and CP13* D70N (green line) at pH5. All data taken at 25°C.

To evaluate whether CP13* populates a partially folded intermediate at equilibrium, we performed proteolysis experiments at both pH 5 and pH 7. Proteolysis of CP13* was monitored as a function of time using the non-specific protease thermolysin^20,21^. Quenched reactions from different time points were run on an SDS-PAGE gel to quantify the remaining full-length protein (Fig 4A and Materials and Methods). There is an exponential decrease in the intensity of full-length protein at both pH 5 and pH 7. Concomitant with this disappearance of the full-length protein, a smaller band appears at approximately 17kDa (I_c_) (Fig. 4A). The size of this proteolysis-resistant fragment is consistent with the size of the C-terminal domain. Mass spectrometry analysis indicates that the proteolysis fragment sequence maps onto the folded region of the NSHX intermediate (I_c_ is the same as I_eq_)^10^. Together, these results suggest that the cleavable state consists of an unfolded N-terminal subdomain and a folded C-terminal domain.

**Figure 4.**
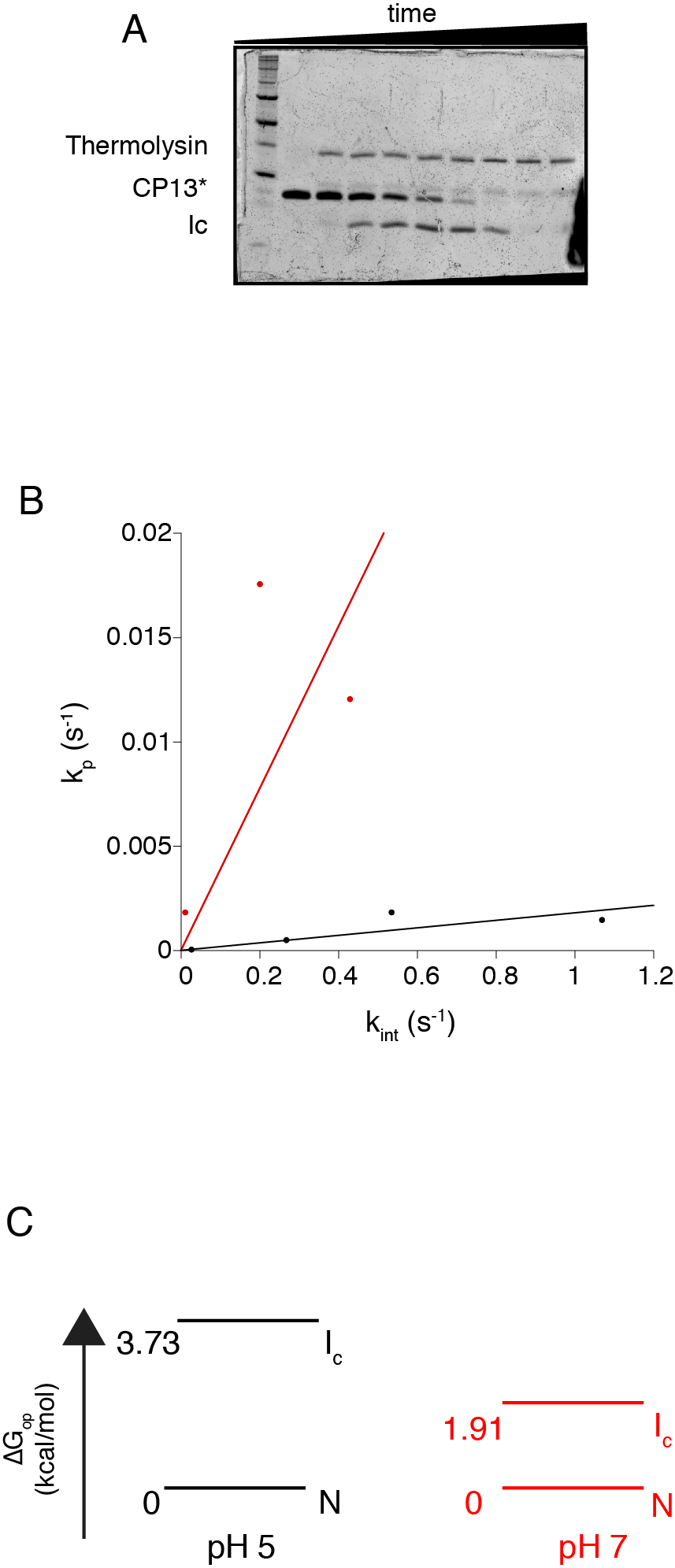
Proteolysis of CP13* and variants. A. Representative gel of CP13* proteolysis. CP13* was incubated at 25°C with 0.20mg/ml of thermolysin at pH 5 or 7 (See Materials and Methods). Samples taken at designated time points and run on a SDS-PAGE gel. Cleavage product denoted by (I_c_). B. Regime test of proteolysis of CP13* at pH5 (black line) and pH7 (red line). k_p_ was calculated from band intensities for full-length CP13* fit to a single exponential equation. kint was calculated based on the concentration of thermolysin at a given pH. C. Boltzmann diagram of CP13* at pH 5 and 7 determined from proteolysis experiments.

To gain more information about the difference in the energetics of CP13* under these two conditions we carried out a quantitative analysis of the observed proteolysis rate (*k*_*p*_), utilizing formalisms developed for hydrogen exchange^21–23^. This approach allows for the determination of either the kinetics or thermodynamics of the opening reaction that leads to the formation of the cleavable conformation, depending on the conditions of the experiment. To determine whether proteolysis in our experiments was in the kinetic (EX1 exchange) regime or thermodynamic (EX2 exchange) regime, we monitored *k*_*p*_ as a function of the intrinsic rate of proteolysis, *k*_*int*_, by modulating the concentration of thermolysin. If *k*_*p*_ is constant as a function of *k*_*int*_, *k*_*p*_ reports on the kinetics of the partial unfolding reaction. If *k*_*p*_ changes linearly as a function of *k*_*int*_, *k*_*p*_ provides thermodynamic information about the reaction, and the slope of this line is equal to the equilibrium constant of the opening reaction (K_*prot*_)^23,24^. As shown in Figure 4B, there is a linear relationship between *k*_*p*_ and *k*_*int*_ for CP13* at both pH 5 and 7. Therefore, we are in the EX2 regime, and *k*_*p*_ reports on the thermodynamics of the unfolding reaction. Using the slopes of these lines (slope_pH5_= 0.0018, slope_pH7_=0.039) we find that ΔG_prot_, the free-energy difference between the ground state (N) and the cleavable conformation (I_c_), is 3.73 kcal/mol at pH 5 and 1. 91 kcal/mol at pH 7. These data suggest that destabilizing the salt bridge, increases the population of the intermediate which explains the difference in m-values under these two conditions.

The pH dependence of the intermediate population implicates a role for the H31-D70 salt bridge in coupling between the two subdomains of CP13*. To probe this directly, we performed equilibrium denaturation experiments with the single-site variants of CP13* H31N and D70N. Even at pH 5, two-state fits of the urea-induced denaturation curves for both CP13* H31N and CP13* D70N result in m-values significantly lower than that for CP13* (CP13* H31N: 2.10 ±0.05 kcal/mol*M (Δm-value= 0.72 kcal/mol*M); CP13*D70N: 2.17 ±0.03 kcal/mol*M (Δm-value= 0.65 kcal/mol*M)) (Table I and Fig. 4). Interestingly, these m-values are quite similar to each other and to the m-value of a protein fragment designed to mimic I_eq_ observed in previous NSHX experiments^9,10,12^ LCTF, 2.06 ±0.02 kcal/mol*M (Table I). These lower m-values, together with the decreased m-value of CP13* at pH 7 compared to pH 5, suggest that the salt bridge (H31-D70) plays an important role in the folding cooperativity of the two subdomains in CP13*, without which there is a significant population of an intermediate. Indeed, proteolysis studies on these two variants show complete proteolysis to I_c_ within one minute (the first time point). This rapid proteolysis suggests that, in these variants, the N-terminal subdomain is substantially unfolded under native conditions.

## Discussion

In this work, we investigate how a helix spanning the two subdomains of T4 lysozyme*, CP13*, couples distant regions of the protein. CP13* is a small globular protein composed of two distinct subdomains with almost no side-chain interactions between the two^7,8^, yet it exhibits typical two-state equilibrium unfolding behavior^6,8–10^, suggesting that the coupled folding of the two subdomains must arise from sources other than interactions at the subdomain interface. We investigate the role of the single topological connection between the subdomains, the C-helix in coupling these subdomains.

By comparing the stability of the N-terminal subdomain in different contexts - in isolation, with additional residues to complete the C-helix, with additional residues encoding a stable helix, and in the context of the full-length protein - we find that the energetics of this subdomain are tunable and that this malleability relies on the addition of a stable helical sequence to the N-terminal subdomain, thus suggesting a role for the C-helix in stabilizing this region. By investigating the pH dependence of the subdomain’s stability and the effect of site-specific mutations, we find evidence that these changes in stability directly correlate with a well-known salt bridge within the N-terminal subdomain^13^. The stability of this interaction increases as the stability of the helix is increased remotely (its C-terminus). In the context of the full-length protein, where interactions between the C-terminal end of the C-helix and other residues of the C-terminal subdomain maximally stabilize the helix, we find that removing the salt bridge decouples the two subdomains such that a conformation with an unfolded N-terminal subdomain is populated at equilibrium. Because D70 is in the C-helix, we posit that stabilizing the helix orients D70 to strengthen the D70-H31 salt bridge. Thus, the stability of the C-helix provides the conduit for long-range energetic coupling of the two subdomains.

### Coupling of the subdomains via the C-helix

Our current work, together with previous studies of the C-terminal subdomain, demonstrates that both subdomains can fold autonomously, as suggested by analyses of side-chain contacts^7^ (see figure 1B for a contact map of CP13*). Independently these regions are marginally stable, but when fused in the context of CP13* the stability is much greater than the sum of the parts (3.7^5^ kcal/mol and 1.60 kcal/mol vs. 12 kcal/mol (Table I)).

In CP13*, the two subdomains are joined only by a shared 23 amino-acid helix, the C-helix. The first sixteen residues are part of the N-terminal subdomain while the last seven are part of the C-terminal subdomain^8^. While helical in the full-length protein, these residues in isolation do not encode a stable helix: a peptide corresponding to the isolated C-helix is unstructured in aqueous solutions when monitored by far-UV CD^25^, and AGADIR, an algorithm parameterized to predict peptide helix propensity, suggests a very low helical propensity for the C-helix sequence (4.3 %)^26^.

The C-helix is stabilized in the context of the C-terminal subdomain. Previously we determined the high-resolution crystal structure of a fragment encoding the C-terminal subdomain with the entire C-helix (an additional 15 residues: 60-74), LCTF^10^. LCTF has a native-like fold and residues 65-81 form a helix, indicating that all but the first turn of the C-helix (residues 59-64) folds in this context. NSHX data shows that C-helix residues 75-81, which make contacts with other residues in the C-terminal subdomain, are significantly more stable than the remainder of the C-helix^10,12^. Thus, the C-terminal subdomain and its contacts with the last seven residues of the C-helix (75-81) stabilize the entire helix^10^. Such contact-assisted helix formation has been observed in molecular dynamic simulations of barnase and Protein A^27^.

The presence of the complete C-helix sequence also appears to stabilize the C-terminal subdomain. Interactions between the C-helix and the rest of the C-terminal subdomain stabilizes that subdomain, as LCTF is 5.5 kcal/mol more stable than CTF (75-164, 1-12)^9,10^. In the present work, we find that the N-terminal subdomain shows similar, albeit much less, stabilization and linkage between the C-helix and the subdomain. The presence of the complete C-helix sequence stabilizes both subdomains in isolation.

### Long-distance coupling and communication via helix stability

In proteins, relatively weak bimolecular interactions are strengthened when they occur between components of a single polypeptide chain due to an increase in effective concentration^28^. For example, the interaction between imidazole and carboxylic acid is quite weak; however, in the case of the H31-D70 salt bridge in T4L*, it is worth 3-5 kcal/mol^13^. The stabilization of this interaction can be thought of as a result of the increased effective concentration of H31 and D70 in the context of the protein.

Here, we observe an increase in the stability of the H31-D70N interaction when the NTF* sequence is extended either by completion of the C-helix (LNTF*), by appending a stable engineered alpha helix (NTF-αH*), or within the context of the full-length protein (CP13*). These data suggest that increasing the stability of the C-terminal end of the C-helix modulates the effective concentration of H31 and D70, presumably by positioning them in an optimal orientation for interaction.

Our model for the role of the C-helix in coupling the N- and C-terminal subdomains is that energetic information is transmitted through helix stabilization. We suggest that side-chain interactions between residues in the C-terminal subdomain stabilize the alpha helix via interactions with the C-terminus of the C-helix. The helix-coil transition is known to be a cooperative process – nucleation or stabilization at one end of the helix, will propagate to the rest of the helix^29,30^. Thus, these side-chain interactions in the C-terminal subdomain stabilize the entire C-helix, including the portion that extends into the N-terminal subdomain. The resulting proper and stable orientation of D70, which resides on the C-helix, increases its effective concentration and its interaction with H31, thereby stabilizing the N-terminal subdomain.

It appears that an important aspect of our model is the low intrinsic stability of the C-helix^25^. This property allows helix stability to be tuned via side-chain interactions in the individual subdomains, which provides the needed coupling to ensure two-state folding. If the C-helix sequence encoded an intrinsically stable helix, this would likely prohibit the communication and coupling between the subdomains. Therefore, it seems likely that the C-helix’s lack of intrinsic helicity in aqueous solution^25,26^ is what allows it to act as a conduit of structural and energetic information in the T4L* sequence and prevent substantial build-up of a partially folded intermediate that may lead to deleterious aggregation or amyloid formation^31^.

Several studies on the nature of coupling between regions of a cooperatively folded protein have focused on side-chain interactions at the interfaces between each region. For example, elegant studies on variants of ankryin-repeat proteins have highlighted the importance of the interface interactions between neighboring repeat elements and their relationship to the intrinsic stability of each unit in creating a cooperatively folded protein^32^. For titin, a multi-subunit protein with many Ig-like β-sheet modules, the domains are coupled by mutually stabilizing interactions between domains ^33^.

Communication via a unit of secondary structure, such as an alpha helix, provides an alternative and important mechanism for coupling systems without notable interfaces or side-chain contacts between the modules. A helix-dependent mechanism of coupling has previously been identified in the model protein spectrin. In nature, spectrin exists as a multidomain protein, with each domain composed of three α-helices. In the multidomain context, a central helix spans two domains. Similar to the subdomains of T4L*, isolated spectrin domains unfold in a two-state manner at equilibrium, but they are more stable in the multidomain context, suggesting a role for the central helix in coupling the energetics of these domains. Despite the similarities between spectrin’s behavior and our results for T4L*, there is a notable distinction. A study of how cooperativity is conferred in pairs of spectrin domains found that the sequence of the central helix is important for proper coupling between spectrin domains^18^. This helix sequence-dependent communication between these domains differs from our model, as it relies on the specific sequence of the linking helix, whereas the necessary feature in our model is a lack of encoded helicity.

### Considerations for protein dissection

A common practice in biochemical studies of large mulitdomain proteins is to isolate a given domain or domains using sequence homology and secondary structure predictions as guides. Often regions with no predicted secondary structure are thought to serve as flexible linkers that play an insignificant role in the folding of neighboring domains, and are most often used as excision sites. The role of the C-helix in the cooperativity of T4L*, in spite of its lack of intrinsic helicity, suggests a need for caution when evaluating seemingly inconsequential regions of protein sequence during protein dissection studies as they might be critical to the stability and folding of neighboring sequences.

### Implications for co-translational folding

In addition to offering a mechanism to explain long-distance communication in fully translated proteins, our helix stabilization model might be relevant to co-translational folding. The ribosomal exit tunnel can accommodate^34–37^ and stabilize helices^38^, but the potential role of these transient helices remains unclear. We posit that the stabilizing effect of the tunnel on these transient helices might be transmitted to the exposed nascent polypeptide chain, thereby stabilizing folding of the emerging N-terminus.

## Materials and Methods

### Plasmid Construction

The NTF* construct was subcloned from the CP13* sequence into a pET27a vector that included a sequence that encodes an N-terminal TEV-cleavable hexahistidine tag. Site-directed mutagenesis was used to convert the TEV-encoding site to a HRV-3C protease cleavable site. The LNTF* construct was constructed from a CP13* construct containing encoded N-terminal hexahisitidine tag and a HRV-3C site by site-directed mutagenesis to include a stop codon after residue 81. NTF-αH* was created using the NTF* construct using site directed mutagenesis to create the alanine-based helix sequence (AEAAAKEAAAKEAAAKA). All variants were made using site-directed mutagenesis.

### Protein Expression and Purification

Full-length proteins were expressed as described previously^9,10^. All fragments were overexpressed in BL21 Codon plus cells. Cells were grown in Luria Broth containing kanamycin (50μg/ml) at 37°C For NTF*, LNTF*, and NTF*-αH protein expression was induced with 1mM IPTG at 0D_600_~0.6. Cells were harvested after 3 hours post induction.

Pellets of cells that overexpressed NTF*, LNTF, NTF*-αH and their variants, were resuspended in 20mM Tris pH 8.0 500mM NaCl 20mM imidazole 6M GdmCl. The resuspended pellets were sonicated and whole cell lysates were spun and filtered. Filtered lysate was run on a column packed with Ni-NTA agarose beads. The protein was eluted from resin using 20mM Tris pH 8.0 500mM NaCl 500mM imidazole and 6 M GdmCl. The eluent of NTF-αH was dialyzed into 20mM KOAc 50mM KCl. The eluents of NTF, LNTF and their variants were concentrated and buffer exchanged into 20mM KOAc 50mM KCl pH 5.0 and 4M urea and concentrated further. Aliquots were dialyzed into denaturant-free buffer to ~1mg/ml prior to experiments.

Cell pellets containing overexpressed CP13*, were resuspended into 20mM Tris pH 8.0 10mM NaCl. After sonication and centrifugation the lysate was run on an S column with a gradient against 20mM Tris pH 8.0 300mM NaCl. The peak fractions were then diluted into 20mM NaOAc 10mM NaCl pH 4.5 and run on an S column with a gradient against 20mM NaOAc 1M sodium chloride pH 4.5. The peak fractions were collected, and dialyzed into 20mM KOAc 50mM KCl pH 5.0.

CP13* D70N and H31N were purified from inclusion bodies. The cells were lysed and centrifuged. Pellets were washed by sonication in 20mM Tris pH 8.0 and 1% Triton X-100 and centrifuged. The wash step was repeated without Triton and centrifuged. The pellet was solubilized by sonication in 20mM Tris pH 8.0 6M GdmCl and centrifuged. The solubilized protein was added dropwise into 20mM Tris 10mM NaCl pH 8.0 and purified over an S column as described above.

### Circular Dichroism Spectra

Far-UV experiments were performed on an Aviv 410 spectrophotometer. All data was taken in either 20mM KOAc pH 5.0 50mM KCl or 20mM KPO4 pH 7.0 50mM KCl. CD spectra were taken in an AVIV 410 CD spectrometer at 4°C in a 0.1 cm quartz cuvette at protein concentrations ~500 μg/ml. Data were collected between 260-200 nm at 1 nm intervals with each data point an average of 5 seconds of data. Data with dynode above 480v were not included.

### Equilibrium Denaturant Melts

All equilibrium experiments were carried out in aforementioned buffer conditions. Urea melts of NTF*, LNTF* and NTF-αH* and variants were carried out at 4°C, those of CP13* and variants were carried out at room temperature. CD signal was monitored at 222 nm and carried out in a 1cm quartz cuvette. Chemical denaturant melts were carried out using an automated titrator with 5-minute equilibration times at each denaturant concentration. Protein concentrations ranged from 20-50 μg/ml.

### Proteolysis

Proteolysis rates were measured by incubating protein (400ug/ml) with various concentrations (0.01mg/ml, 0.1mg/ml, 0.2mg/ml, and 0.4 mg/ml) of thermolysin at 25°C at pH 5 or 7 using the buffers described above. Proteolysis reactions were quenched at various time points with 50mM EDTA. The quenched reactions were run on SDS-PAGE gels. Gels were stained with SyproRed (Lonza Rockland) and scanned using a Typhoon scanner (GE Healthcare). ImageJ (NIH) was used to quantify the band corresponding to the full-length protein. The band intensities were plotted as a function of time in Kaleidagraph (Synergy Software) and fit to a first-order rate equation to calculate k_p_. The k_p_ at various thermolysin concentrations were plotted against k_int_ which was calculated from the k_cat_/K_m_ values calculated previously. The data were fit to a line with a fixed resulting line was fit and the slope of that line (which is K_op_) was used to calculate ΔG_prot_ (= −RTlnK_prot_)

## Acknowledgements

This work used the Vincent J. Proteomics/Mass Spectrometry Laboratory at UC Berkeley, supported in part by NIH S10 Instrumentation Grant S10RR025622. We thank the entire Marqusee lab, Rachel Bernstein, Laura Rosen, and Katherine Tripp for helpful comments and discussion. This work was supported by a grant from the NIH (R01-GM050945).

